# Next-generation sequencing of double stranded RNA is greatly improved by treatment with the inexpensive denaturing reagent DMSO

**DOI:** 10.1101/644591

**Authors:** Alexander H. Wilcox, Eric Delwart, Samuel L. Díaz Muñoz

**Author notes:** Address correspondence to: Samuel L. Díaz Muñoz.

## Abstract

Double stranded RNA (dsRNA) is the genetic material of important viruses and a key component of RNA interference-based immunity in eukaryotes. Previous studies have noted difficulties in determining the sequence of dsRNA molecules that have affected studies of immune function and estimates of viral diversity in nature. Dimethyl sulfoxide (DMSO) has been used to denature dsRNA prior to the reverse transcription stage to improve RT-PCR and Sanger sequencing. We systematically tested the utility of DMSO to improve sequencing yield of a dsRNA virus (Φ6) in a short-read next generation sequencing platform. DMSO treatment improved sequencing read recovery by over two orders of magnitude, even when RNA and cDNA concentrations were below the limit of detection. We also tested the effects of DMSO on a mock eukaryotic viral community and found that dsRNA virus reads increased with DMSO treatment. Furthermore, we provide evidence that DMSO treatment does not adversely affect recovery of reads from a single-stranded RNA viral genome (Influenza A/California/07/2009). We suggest that up to 50% DMSO treatment be used prior to cDNA synthesis when samples of interest are composed of or may contain dsRNA.

**Data Summary:** Sequence data was deposited in the NCBI Short Read Archive (accession numbers: PRJNA527100, PRJNA527101, PRJNA527098). Data and code for analysis is available on GitHub (https://github.com/awilcox83/dsRNA-sequencing/, <u>doi:10.5281/zenodo.1453423</u>). Protocol for dsRNA sequencing is posted on protocols.io (<u>doi:10.17504/protocols.io.ugnetve</u>).

## Introduction

Ribonucleic acid (RNA) is a ubiquitous biological molecule involved in transcription and translation, which also serves as the genetic material of a large number of important viruses. The double-stranded form of RNA (dsRNA) is believed to be less abundant in nature, but is a crucial component of a number of biological systems. It has a central role in the RNA interference system [1], which modulates innate immunity in plants and animals, and serves as a replicative intermediate of (+) ssRNA viruses, while also being present in dsDNA and (-) ssRNA infections [25]. Moreover, it also serves as the genetic material of a number of virus lineages (including the families *Reoviridae, Cystoviridae, Picobirnaviridae*) that infect humans, animals, plants, fungi, and bacteria; which play important medical, ecological, and scientific roles. A number of dsRNA viruses are of clinical and agricultural significance, such as Bluetongue virus, which causes high morbidity and mortality in ruminants [2], and rotavirus, which causes acute gastroenteritis in humans [3]. There are indications that the overall diversity of RNA viruses may be underestimated [4, 5] and difficulties sequencing dsRNA in particular have been noted in the literature [6,7]. As a consequence, dsRNA virus lineages may be underrepresented and dsRNAs involved in immunity may be underestimated.

## Theory and implementation

The extent of microbial diversity has been revealed by powerful whole genome sequencing tools that are quickly becoming standard tools in biology. However, next-generation sequencing has known biases according to the nucleic acid composition [8]. A major limitation is that most sequencing platforms cannot sequence RNA directly, requiring that it is first reverse-transcribed into its complementary DNA (cDNA). cDNA synthesis is typically achieved by hybridising oligonucleotide DNA primers to the RNA and using a reverse transcriptase to synthesise the remainder of the complementary DNA strand. This step poses a particular problem to dsRNA because the presence of a complementary strand blocks the ability of these primers to bind. The blocking has a direct effect on the amount of dsRNA converted to cDNA, resulting in many fewer sequencing reads being generated relative to the true amount of RNA present. Additionally, many dsRNA viruses have small genomes that limit the amount of RNA that can be extracted, further complicating the determination of dsRNA sequences.

As early as 1968, dimethyl sulfoxide (DMSO) has been shown to have a denaturing effect on nucleic acids [9]. DMSO has been successfully used to improve the performance of RT-PCR [10] and Sanger Sequencing [11]. However, to our knowledge DMSO has not been used for next-generation sequencing approaches and many dsRNA sequencing studies omit DMSO treatment [6, 12, 13, 14]. Moreover, there is no standard protocol for DMSO treatment of samples and previous methods vary greatly in their conditions, particularly DMSO concentration, which has ranged from 15% [11] to 90% [15].

This paper investigated four questions regarding the effect of DMSO treatment on next-generation sequencing: 1) Does DMSO treatment improve recovery of dsRNA reads and at what concentration? 2) Does DMSO affect read coverage and accuracy of a viral genome? 3) Is the effect of DMSO independent of RNA concentration? 4) Does DMSO treatment negatively affect the recovery of single-stranded RNA (ssRNA) genomes? Our results suggest that treatment with a high concentration of DMSO greatly increases the number of reads generated when sequencing dsRNA with no effect on read accuracy, without adversely affecting sequencing of ssRNA virus genomes on the Illumina short-read sequencing platform.

We carried out the methods in this paper on two different viruses: *Pseudomonas* phage Φ6, which has a dsRNA genome made up of three segments, and human influenza virus A H1N1, which contains eight ssRNA segments. We also used a mock eukaryotic viral community, manufactured by the National Institute for Biological Standards and Control (NIBSC, UK) as a reference material for multiplex viral detection (NIBSC reagent 11/242-001). This reagent was expected to contain 25 human pathogenic viruses and has been used to investigate viral detection methods on mixed and metagenomic samples [16].

### Sample Preparation

Φ6 lysate was prepared by plating phage and its *Pseudomonas syringae* host using double agar overlay. Phages were harvested by selecting a plate with semi-confluent lysis, transferring the soft agar layer to 3ml LB (Lennox) media, and centrifuging to remove host cells and agar. An influenza virus lysate was generated from egg-passaged stock of influenza A/California/07/2009(H1NI) (generously provided by Ted M. Ross, University of Georgia), which was expanded by passage at a low multiplicity of infection in Madin-Darby canine kidney (MDCK) cells in culture. Viral lysates and the mock viral community (NIBSC reagent 11/242-001) were passed through a 0.22μm millipore polyethersulfone membrane filter (Millex) to remove debris and contaminants. 1ml of each filtrate was treated with 25μl DNAse I (Thermo Scientific) and 50μl RNAse A/T1 mix (Thermo Scientific) with 1X DNAse I Buffer (Thermo Scientific) at 37°C for 1 hour 30 minutes to degrade extracapsular nucleic acids. Viral RNA was extracted using an RNeasy Mini Kit (Qiagen), passing a total of 900μl of nuclease-treated lysate through a column, and eluting into 100μl of elution buffer.

### DMSO Treatment, reverse transcription and sequencing

Viral RNA samples were divided into 20μl aliquots. DMSO was added to concentrations (v/v) of 15%, 50% and 90% for each sample, followed by 1 hour and 30 minutes of incubation at 65°C. DMSO was removed using a RNeasy MinElute Cleanup Kit (Qiagen, Valencia CA), following the manufacturer’s instructions. An additional sample was treated by heat denaturation but not DMSO: the tube containing the RNA extraction was placed in boiling water for five minutes [27]. Following this, all samples were placed on ice until cDNA synthesis was carried out. Other than heat or DMSO treatment, all samples followed standard cDNA synthesis methods.

First strand cDNA synthesis was carried out as described in the SuperScript III First Strand cDNA Synthesis kit (Fisher) instructions, by adding 5μl of each RNA sample (including a control that had not undergone DMSO treatment or column cleanup) to 1μl of random hexamer oligos, 1μl dNTPs and 3μl DEPC-treated water. Reactions were incubated at 65°C for 5 minutes then placed on ice for 1 minute. 1X reverse transcriptase buffer, 5mM McCl_2_, 0.01M DTT, 1μl RNAseOUT and 1μl Superscript III RT enzyme were added to each reaction for a total volume of 20μl. Reactions were incubated on a thermal cycler at 25°C for 10 minutes, 50°C for 50 minutes and 85°C for 5 minutes. Second strand synthesis was carried out by adding 1μl dNTPs, 0.5μl DNA ligase, 2μl DNA polymerase I, 0.5μl RNAse H in 1X second strand synthesis buffer, and made up to a total volume of 40μl with nuclease free water. Reactions were incubated at 16°C for 5 hours and cDNA was purified with a Nucleospin Gel and PCR Clean-up kit (Macherey-Nagal, Düren Germany).

Libraries were prepared for Illumina sequencing using the Nextera XT DNA Sample Preparation Kit (Illumina, San Diego CA), with a 1/5 “scaled” library preparation protocol after Baym *et al*. [17].

### Bioinformatics

Reads were trimmed for adapters and quality using Cutadapt [18] and Sickle [19]. Due to a short fragment size, reads overlapped and so paired-end libraries were merged into single-end libraries using PEAR [20]. For phi6 and influenza virus lysates, these libraries were mapped to the reference genomes using Bowtie2 [21], and bam-readcount [22] was used to determine the read depth at each position. Plots were generated using the ggplot2 package in R.

For the mock viral community, a custom virus discovery pipeline was used to analyze sequencing reads [23]. Reads were translated and aligned to a viral proteome database (consisting of all annotated full or near full viral genomes) using BLASTx. The significant hits to the virus database were then aligned to a non-virus-non-redundant (NVNR) universal proteome database using BLASTx. Hits with more significant E-value to NVNR than to the virus database were removed.

To test if DMSO treatment had any effect on sequencing fidelity we used two approaches to estimate sequencing error rates. First, we used freebayes [24] to generate a VCF file containing all differences from the reference genome with frequency of <5% and Phred quality score of at least 30. A custom Python script was used to count the number of these mutations and the total bases sequenced for each sample, and to calculate the true error rate. We tested for statistically significant differences in error rates among DMSO treatments using a proportion test. Because reference-based approaches to error estimation face limitations, we implemented a reference-free approach to error estimation [26]. We calculated the error rate for each of our DMSO treatments, as implemented in the R package ShadowRegression, and tested for differences using robust linear regressions.

## Results and Discussion

Our findings show that DMSO treatment has a dramatic effect on dsRNA sequencing (Table 1). When we prepared viral lysates for sequencing without DMSO, we did not obtain sufficient reads to cover the entire Φ6 genome. Treatment with 15% DMSO increased the number of mapping reads over sevenfold, whilst the 50% and 90% treatments increased the number of reads by over two orders of magnitude, allowing the full genome to be sequenced at high coverage (average read depth of 1727 for 50% DMSO and 1493 for 90% DMSO).

**Table 1.** The total number of reads generated during sequencing of influenza and Φ6 after treatment at varying DMSO concentrations, and the number of those reads that mapped to reference genomes.

| DMSO % | Total reads | Reads that map to phi6 | Reads that map (%) |
| --- | --- | --- | --- |
| 0 | 1551220 | 1377 | 0.1% |
| 15 | 1314520 | 10358 | 0.8% |
| 50 | 1077133 | 263993 | 24.5% |
| 90 | 1393569 | 224562 | 16.1% |

These increases in genome coverage occurred despite very low starting nucleic acid concentrations. We used a Qubit RNA High Sensitivity Assay kit to quantify RNA immediately after extraction; in all cases RNA was undetectable and so assumed to be under the kit’s limit of detection of 5ng/μl. Additionally, we used a Qubit DNA High Sensitivity Assay kit to quantify the amount of cDNA synthesised. Despite this kit’s lower limit of detection of 200pg/μl, DNA was still not detected. Thus, for dsRNA the raw quantity of starting material may not be as important as the efficiency of cDNA synthesis, a fact that should accounted for when preparing quality control thresholds before next generation sequencing.

We next sought to determine whether DMSO treatment affected other types of RNA present in the sample, which would occur in metagenomes, clinical samples, or transcriptomes. DMSO treatment did not appear to affect the recovery of ssRNA-derived reads from the influenza virus genome. There was no discernible effect when the DMSO concentration was varied (Table 1). The number of reads mapping to the reference genome did decrease from the 0% DMSO treatment to the 15% DMSO treatment. However, this loss in mapping reads was most likely caused by the extra column cleanup step required to remove the DMSO (note that the total reads also decreased). We note that this decrease in reads in the influenza virus sequencing was less than one order of magnitude and still resulted in extremely high read depth, with an average of over 1000x for every influenza segment after DMSO treatment (Table 2). This effect was obscured in Φ6 due to the large increase in mapping reads from DMSO treatment.

**Table 2.** The average read depth for each influenza segment under varying concentrations of DMSO.

| DMSO % | Total reads | Reads that map to flu | Reads that map (%) |
| --- | --- | --- | --- |
| 0 | 1614418 | 931915 | 57.7% |
| 15 | 1369886 | 353752 | 25.8% |
| 50 | 1510801 | 295244 | 19.5% |
| 90 | 1257059 | 446549 | 35.5% |

In order to determine if the presence of DMSO affected any other properties of the RNA when used for downstream sequencing, we plotted the coverage at every nucleotide position for each DMSO concentration used. The plot for influenza (Figure 1) showed a distinct, repeating pattern for each segment, with DMSO concentration appearing to have no effect on relative coverage. This indicates that there is no bias in which reads are affected by the DMSO treatment, and that the reads generated are still representative. A similar pattern could be observed for the 50% and 90% DMSO treatments of Φ6 (Figure 2, the low number of reads made this pattern harder to discern in the 15% treatment). Therefore, DMSO treatment did not adversely affect genome-wide coverage patterns of dsRNA or ssRNA viruses.

**Figure 1.**
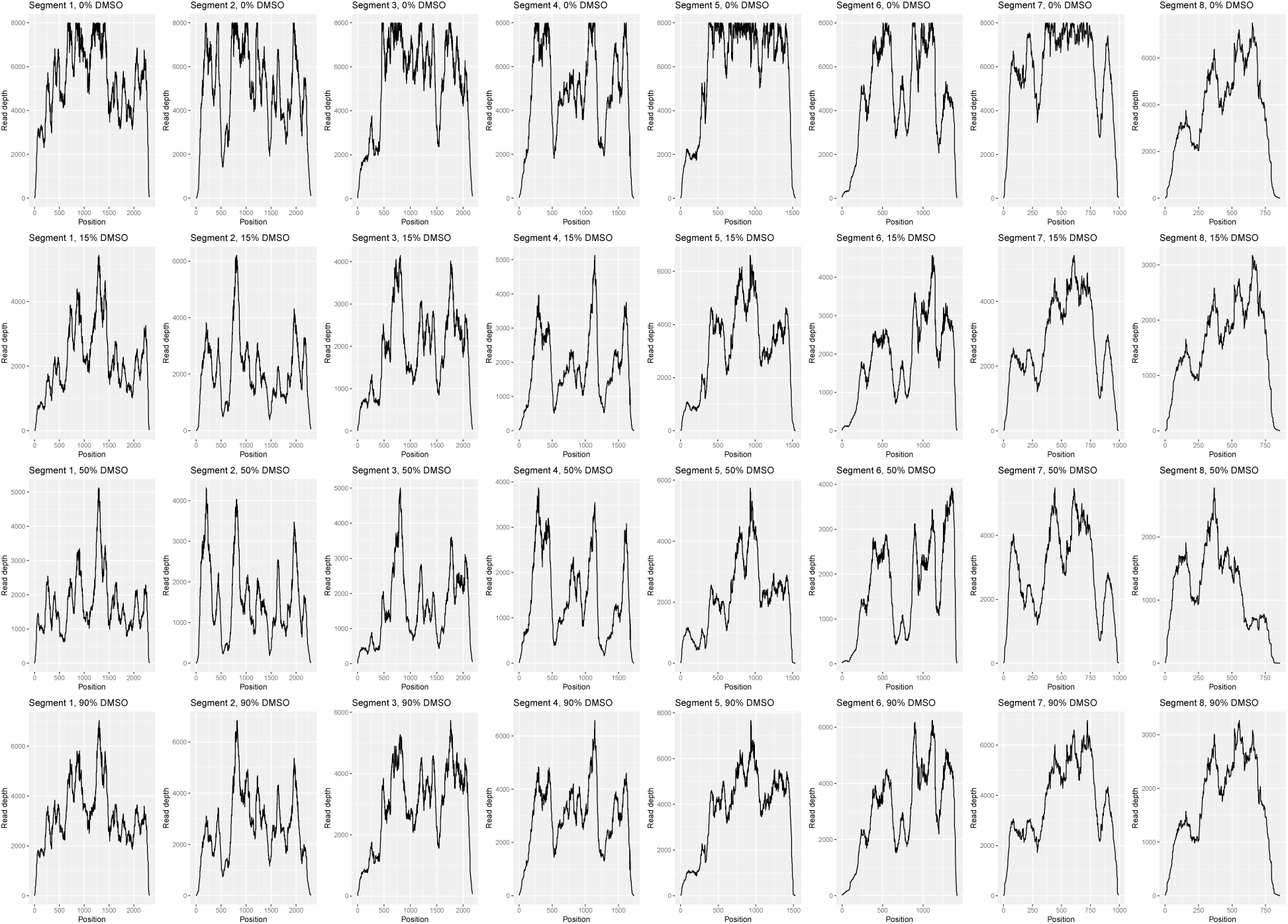
DMSO treatment does not affect sequencing read coverage across the ssRNA influenza genome. Read depth at each position in the influenza genome under varying concentrations of DMSO. Note that 8000 is the maximum read depth supported by the SAM/BAM file format, so some peaks in the 0% DMSO plots have been truncated.

**Figure 2.**
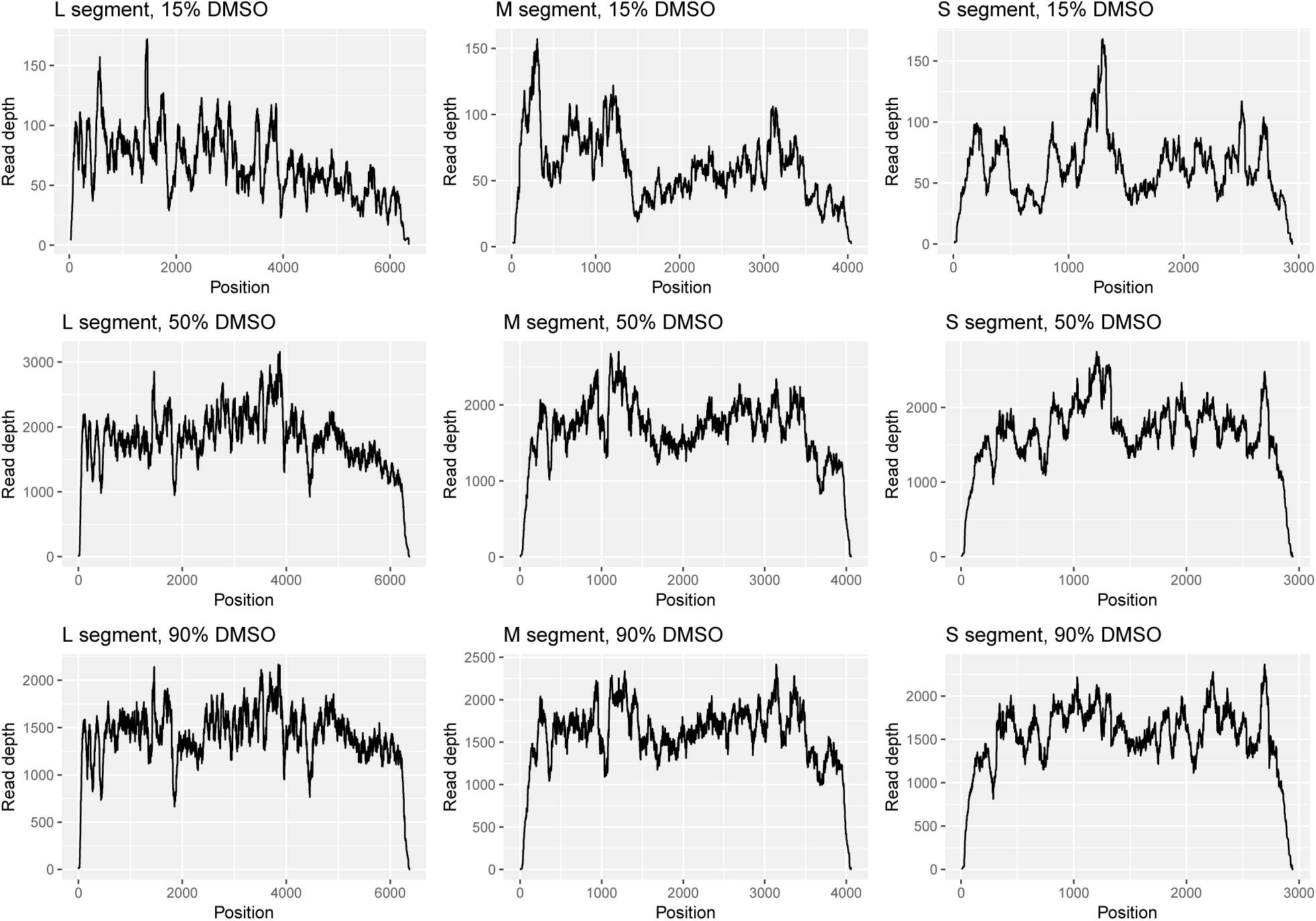
DMSO treatment greatly increases sequence coverage of the dsRNA Φ6 genome. Read depth at each position in the Φ6 genome under varying concentrations of DMSO. There were insufficient reads to generate a plot for 0% DMSO. Read depth at each position in the influenza genome under varying concentrations of DMSO. Note that 8000 is the maximum read depth supported by the SAM/BAM file format, so some peaks in the 0% DMSO plots have been truncated.

To determine if this method worked with higher starting concentrations of RNA, we used an Amicon centrifugal filter unit to concentrate approximately 10ml of Φ6 lysate into 50μl. The concentrated lysate contained 37.6ng/μl RNA (measured by Qubit) and was prepared for sequencing using 90% DMSO (as above), as well as a control without DMSO (heat-treated for 90 minutes at 65°C). While only 0.08% of reads mapped to the reference genome in the non-DMSO treated control, there was an increase to 72.45% in the DMSO-treated sample, again demonstrating the importance of the DMSO treatment over raw RNA concentration (Figure 3). The concentrated sample had a much higher proportion of reads than the non-concentrated sample (16.1%, Table 1), most likely due to the concentration step increasing the ratio of viral RNA to extracapsular RNA. Therefore, DMSO treatment works at varying RNA concentrations and is likely to improve any dsRNA sequencing regardless of starting concentration.

**Figure 3.**
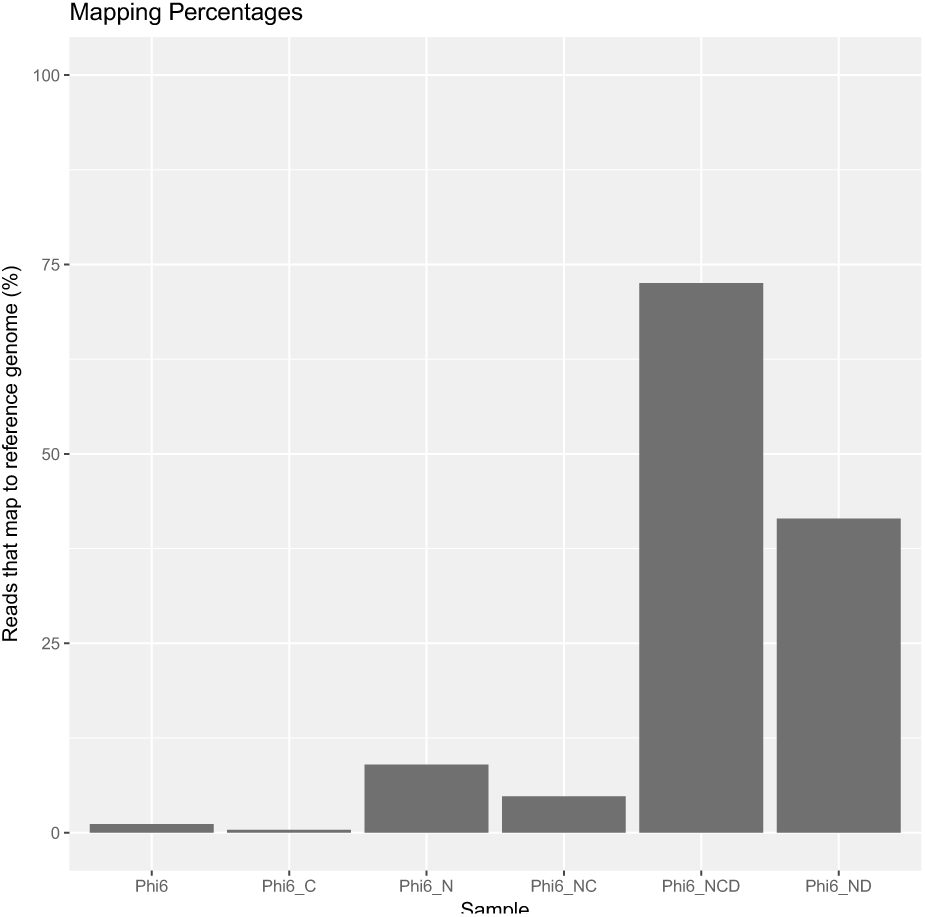
DMSO treatment has a greater effect than RNA concentration on generating dsRNA reads using next generation sequencing. Percent of sequencing reads mapping to phi6 reference genome with no treatment (Phi6), nuclease treatment (Phi6_N), nuclease treatment and concentration (Phi6_NC), nuclease and DMSO treatments and concentration (Phi6_NCD), and nuclease and DMSO treatments.

We also tested if DMSO treatment was more effective than simple heat denaturation. We extracted RNA from Φ6 and divided it into two aliquots. One of these was treated with 50% DMSO as described above, while the other was placed in boiling water for five minutes. After cDNA synthesis, library preparation and sequencing, we mapped the resulting reads to the Φ6 reference genome. In the heat-treated sample, 1.72% of reads successfully mapped to the reference, whilst in the DMSO treated sample this increased to 40.24%. While heat denaturation is clearly of some benefit (compared to 0.08% of reads mapped in the no-treatment control), these data demonstrate that DMSO treatment is the superior method and might be vital when working with low starting RNA concentrations.

Because the fidelity of reverse transcriptase reactions relies on base-pairing, we examined if DMSO treatment had any effect on fidelity by searching for errors in the sequence data using two approaches. First, we extracted all bases that were different from the reference genome with Phred quality score of at least 30. This per-base quality score is equivalent to an expected error rate of 0.1%, meaning over the entire genome sequence the expectation would be 1/1,000 bases to be sequencing errors. Under the assumption that all differences from the reference were errors, these data were used to calculate the true error rate. This error rate was converted to a Phred score (Table 3). If DMSO increased the sequencing error rate, we would expect our calculated Phred score to decrease as DMSO concentration is increased. There was a statistically significant difference in error rates for each DMSO treatment of Φ6 (X_2_ = 533.6, df = 2, p < 2.2e-16), however the error rate did not increase with increasing DMSO concentration (the untreated, control Φ6 sample did not have enough reads for comparison). Similarly, for influenza, the difference in error rates was statistically significant (X_2_ = 575.08, df = 3, p < 2.2e-16), but numerically miniscule (Table 3). Moreover, the error rate decreased as the DMSO concentration increased. In all cases, the true error rate was lower than the expected error rate. Additionally, our calculated error rate is conservative, because of the assumption that all differences from the reference were erroneous. In actuality, some of these differences at low frequency could be true mutations in RNA virus populations (i.e. minor variants).

**Table 3.** The error rate per nucleotide and quality score for each of the Φ6 and influenza samples treated with DMSO.

| Organism | DMSO % | Total errors | Total bases sequenced | Error rate (%) | Phred quality score |
| --- | --- | --- | --- | --- | --- |
| $\Phi 6$ | 15 | 176 | 857974 | 0.021 | 36.9 |
| $\Phi 6$ | 50 | 8940 | 23127006 | 0.039 | 34.1 |
| $\Phi 6$ | 90 | 5276 | 19996871 | 0.026 | 35.8 |
| Influenza | 0 | 34527 | 68561988 | 0.050 | 33.0 |
| Influenza | 15 | 13181 | 29087984 | 0.045 | 33.4 |
| Influenza | 50 | 9549 | 23120765 | 0.041 | 33.8 |
| Influenza | 90 | 16757 | 40369161 | 0.042 | 33.8 |

Second, to estimate error rates in short reads without a reference genome, we also used the ShadowRegression R package [26], which compares reads against each other. Our data shows that DMSO has little effect on the error rate of the ssRNA influenza virus (Figure 4). Using a robust F-test, we found that the difference in influenza error rate between each DMSO-treated sample and the untreated sample was not statistically significant (15%: F = 1.4003, p=0.134; 50%: F = 2.246, p=0.2367; 90%: F = 0.2093, p=0.6473). Moreover, these error rates did not show a pattern with increasing DMSO concentration (Figure 4A, inset). However, in Φ6 we found that as the concentration of DMSO increased, the sequencing error rate actually decreased (Figure 4B, inset). This decrease compared to the 15% DMSO treatment was statistically significant (50%: F = 280.95, p < 2.2×10^−16^; 90%: F = 37.391, p=1.092×10^−9^). Thus, using two statistical approaches, these data collectively indicate that DMSO did not have an adverse effect on read accuracy and in some instances may improve sequencing accuracy.

**Figure 4.**
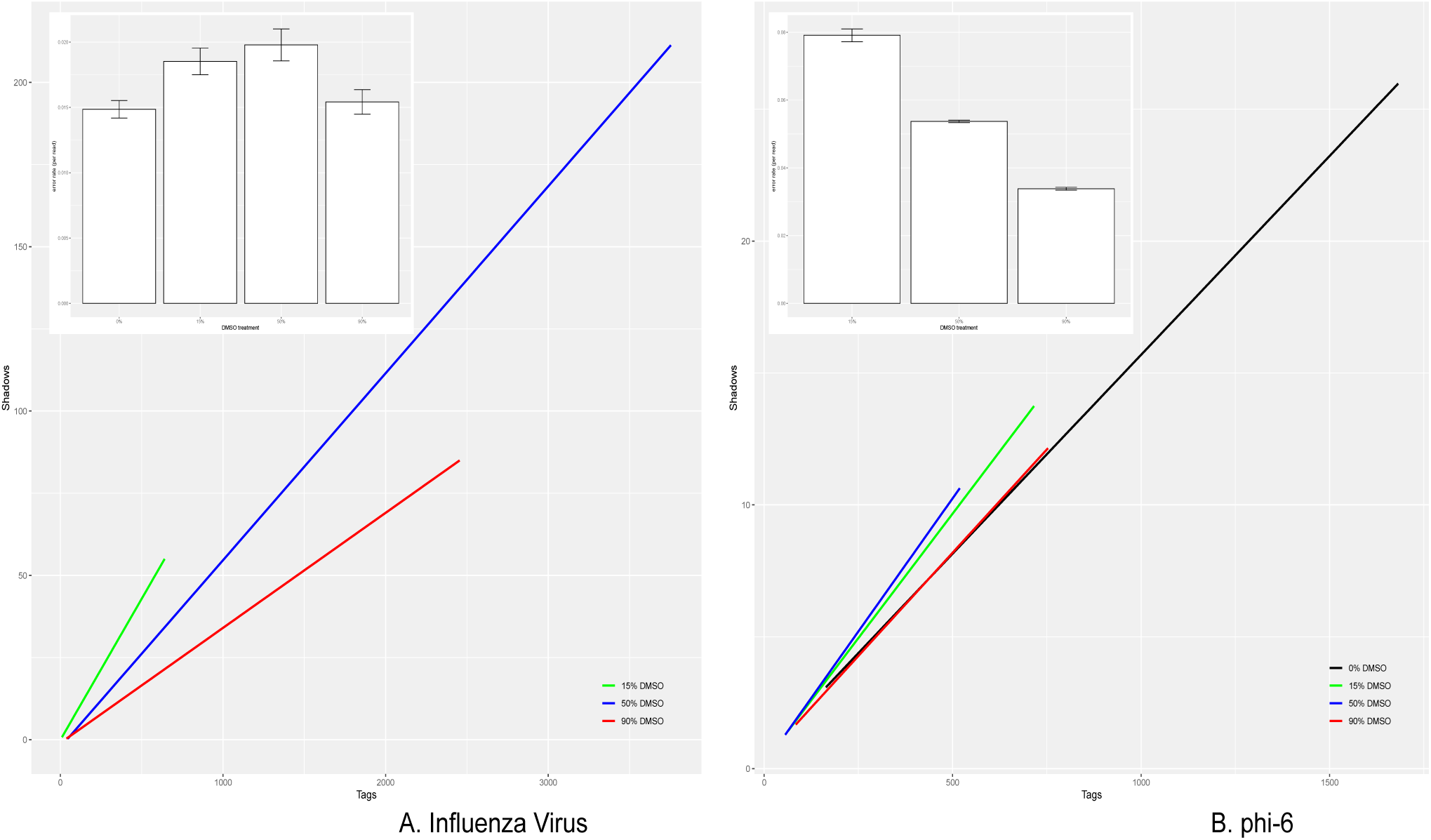
DMSO treatment does not adversely affect sequencing error rates in influenza (A) or phi-6 (B). The R-package ShadowRegression estimates reference-free error rates (inset) based on a transform of the slope of read counts and their “shadows” (main plot line graphs).

The mock viral community (NIBSC reagent 11/242-001) was made by mixing 25 eukaryotic viruses [16]. Of those with RNA genomes, the most abundant in the sample is thought to be the dsRNA rotavirus, based on real time PCR results [23], which we selected for comparison with the ssRNA virus human parechovirus 3. Using a virus discovery pipeline, we only detected 1 read from a dsRNA virus (rotavirus A) in the sample untreated with DMSO (Table 4). However, under the 15% treatment, this increased to over 203 rotavirus reads, as well as reads from dsRNA viruses not thought to have been in the reagent, including human picobirnavirus and some totiviruses. This result is notable because the mock viral community was made such that rotavirus A was the most abundant virus [23]. If the most abundant virus can be so underrepresented in a known sample, this suggests many metagenomic studies will miss almost all dsRNA viruses. Furthermore, we indeed find dsRNA viruses that had previously evaded detection in the mock viral community [16]. The 50% and 90% treatments contained significantly fewer dsRNA reads than the 15% treatment, but reads from all viruses were reduced in these samples. Although the number of reads detected in the DMSO treated samples were low compared with the untreated sample, dsRNA was practically undetectable in the latter. Therefore, despite some caveats we suggest that the power of this method for detecting dsRNA viruses in metagenomic samples is clear.

**Table 4.**
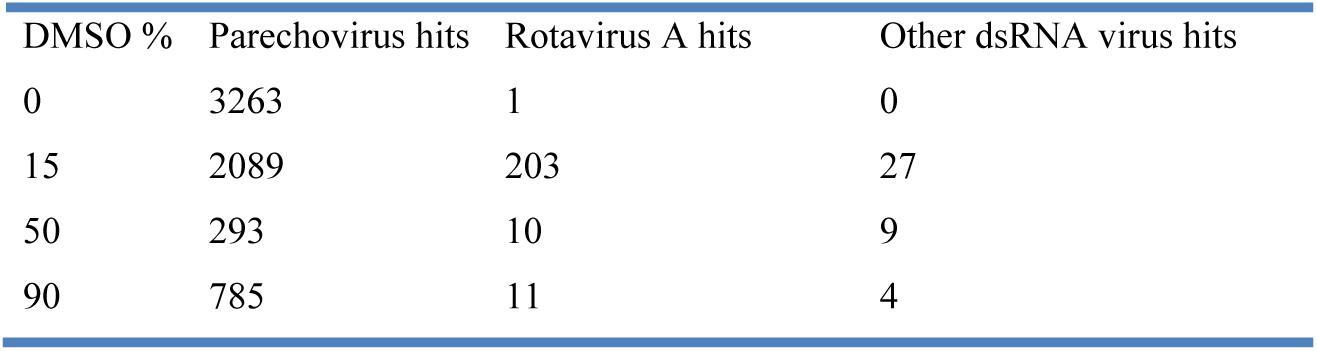
Total number of reads detected in the mock viral community (reagent 11/242-001) under each DMSO treatment for human parechovirus, rotavirus A, and other dsRNA viruses.

## Conclusion

Sequencing of dsRNA can frequently be problematic, with traditional cDNA synthesis being highly inefficient. We have demonstrated that a simple treatment with a cheap and common laboratory reagent can increase the number of sequencing reads from dsRNA organisms by over two orders of magnitude. Importantly, the positive effect of the DMSO treatment occurred independent of RNA concentration, even when RNA was undetectable. Furthermore, DMSO treatment was more important than RNA concentration in determining dsRNA read yield and it did not affect viral genome coverage. We suggest that samples to be sequenced that contain or are suspected to contain dsRNA are treated with at least 50% DMSO prior to cDNA synthesis. This treatment should also improve sequencing of dsRNA involved in innate immunity in plants and animals. We suspect this treatment can be successfully applied to other DNA sequencing technologies, because the DMSO treatment occurs at the cDNA synthesis step and has been shown to improve other procedures such as Sanger sequencing [11].

When preparing an environmental sample for sequencing, it is possible that there may be dsRNA viruses present that are undetectable when following standard protocols. Previous data have shown dsRNA viruses to be underrepresented in metagenomic samples [23]. Our data on the mock viral community (where the putatively most abundant virus was not detected without DMSO treatment) suggests dsRNA viruses will almost invariably go undetected in environmental samples. We have shown that not only will DMSO treatment increase representation of these organisms, the effect on ssRNA representation is minor. It may well be that dsRNA viruses are more numerous than thought, but remain undetected using traditional methods.

## Authors and contributions

Conceptualisation, A.H.W and S.L.D.M.; Methodology, A.H.W.; Software, A.H.W, E.D, and S.L.D.M.; Formal Analysis, A.H.W. and E.D.; Investigation, A.H.W.; Resources, E.D. and S.L.D.M.; Data Curation, A.H.W.; Writing – Original Draft Preparation, A.H.W.; Writing – Review and Editing, and S.L.D.M.; Supervision, S.L.D.M.; Project Administration, S.L.D.M.; Funding, S.L.D.M.

